# Predictive microbial community changes across a temperature gradient

**DOI:** 10.1101/2023.07.28.550899

**Authors:** Xin Sun, Jacquelyn Folmar, Ariel Favier, Nora Pyenson, Alvaro Sanchez, María Rebolleda-Gomez

## Abstract

A central challenge in community ecology is predicting the effects of abiotic factors on community assembly. In particular, microbial communities play a central role in the ecosystem, but we do not understand how changing factors like temperature are going to affect community composition or function. One of the challenges is that we do not understand the mechanistic impacts of temperature on different metabolic strategies, nor how this metabolic plasticity could impact microbial interactions. Dissecting the contribution of environmental factors on microbial interactions in natural ecosystems is hindered by our understanding of microbial physiology and our ability to disentangle interactions from sequencing data. Studying the self-assembly of multiple communities in synthetic environments, here we are able to predict changes in microbial community composition based on metabolic responses of each functional group along a temperature gradient. This research highlights the importance of metabolic plasticity and metabolic trade-offs in predicting species interactions and community dynamics across abiotic gradients.

## Introduction

We currently live in times of rapid change and unprecedented anthropogenic disturbances, making it paramount to understand and predict microbial changes in response to external stressors. Microbial communities break down organic matter and engage in multitrophic interactions (Solden et al. 2018; Pontrelli et al. 2022; Glassman et al. 2018), directly impacting biogeochemical cycles and primary productivity (Nguyen et al. 2022; Kim et al. 2023; Schlesinger and Andrews 2000). Changes in microbial communities’ composition can drastically alter their function, leading to consequences at the ecosystem level (Glassman et al. 2018; Cavigelli and Robertson 2000; Reed and Martiny 2013; Matulich and Martiny 2015).

Forecasting changes in biological communities is a central goal in ecology, yet the complexity of these communities makes this a challenging task. Changes in the environment affect the physiology of individual organisms and interactions between them, leading to changes in the composition and function of biological communities (Dell, Pawar, and Savage 2014; Burnside et al. 2014; Crocker et al. 2023; D. J. Smith and Amarasekare 2018; Amarasekare 2015). These changes in communities generate feedback loops between the biological systems and their abiotic environment, resulting in further shifts in both systems (Suding, Gross, and Houseman 2004; García et al. 2023; Tseng, Bernhardt, and Chila 2019). One way of overcoming these challenges is through the integration of organismal physiology in predicting population dynamics (Brown et al. 2004; Savage et al. 2004).

Temperature is one of the most important abiotic factors structuring past and present ecological communities and their interactions (e.g. Swain et al, this special section). In particular, increased temperatures have a large impact on microbial communities across different ecosystems (Thomas et al. 2012; Wu et al. 2022). Temperature changes affect bacterial physiology (Price and Sowers 2004; Allison et al. 2018; Postmus et al. 2008), species interactions (Amarasekare 2015; Dziallas and Grossart 2012), and therefore community function (Allison, Wallenstein, and Bradford 2010; Xiao et al. 2023; Hall and Cotner 2007). Warming promotes the growth of heterotrophic microbes and accelerates their metabolic rates, which may result in a larger CO_2_ flux— creating a positive feedback between warming and elevated CO_2_ (Davidson and Janssens 2006; Frey et al. 2013). However, the effects of temperature on carbon fluxes depend on (among other things) microbial physiology and community composition (Allison, Wallenstein, and Bradford 2010; Auffret et al. 2016). Thus, it remains unclear how community structure and function will change with increasing temperature. To better forecast (and potentially intervene) we require a better integration of cellular physiology in our understanding of ecological dynamics.

Recent work in microbial ecology has shown that similar environments favor similar functional groups, despite substantial variation at lower taxonomic levels (Louca et al. 2018; Goldford et al. 2018). The simultaneous observation of functional convergence and taxonomic divergence has been explained using synthetic communities, but the impact of different abiotic factors on community assembly remains poorly understood. Replicated community assembly in synthetic habitats with a single limiting carbon source led to reproducible assemblies at the family level (Goldford et al. 2018). These assemblies vary across carbon sources, but similar carbon sources result in similar communities (Estrela et al. 2021). In particular, community assembly in sugars can be described by simple metabolic rules.

Structure in these communities results from a couple of metabolic trade-offs. First, heterotrophic bacteria tend to specialize along an axis of sugar—organic acid preference (Gralka, Pollak, and Cordero 2022). The dominant families in these communities differ in their specialization: fermenter taxa (often of the Enterobacteriaceae family) prefer sugars, whereas respirators (members of Pseudomonadaceae and Moraxelaceae families) grow better in organic acids (Estrela et al. 2022). This specialization makes sense because there is a cost of switching between tasks (Goldsby et al. 2012); there is a trade-off between growth at one of these carbon sources and the time it takes to switch to the other one (Basan et al. 2020). The second trade-off is between fermenters’ growth rate and yield. Respirofermentative bacteria can both ferment or respire sugars. As their growth rate increases, they start fermenting more, decreasing their ATP yield and secreting more organic acids into the media (Basan et al. 2015; Thomas Pfeiffer and Bonhoeffer 2002). These two trade-offs are enough to quantitatively predict the ratio of respirator to fermenter taxa (R/F) in these communities (Estrela et al. 2022).

The predictability of community function in these communities and the ability to monitor and control ecological processes in their synthetic habitats make them an ideal model system to evaluate the impact of different abiotic factors on species interactions and community assembly (Estrela, Sánchez, and Rebolleda-Gómez 2021; Sun and Sanchez 2023). In particular, temperature is rapidly changing and it is an environmental factor central to determining microbial community structure and function (Abreu et al. 2023; Sheik et al. 2011; DeAngelis et al. 2015), but its multifaceted impacts on microbial physiology and interactions make it challenging to predict its consequences on microbial communities.

We hypothesized that as temperature increases we would see an increase in growth rate at the expense of biomass formation, as respiro-fermentative bacteria switch from a more carbon-efficient but slower-growing respirative metabolism to a faster-growing but inefficient fermentatative strategy (T. Pfeiffer, Schuster, and Bonhoeffer 2001). As a consequence, we would expect more organic acid secretions favoring the growth of respirators (Estrela et al. 2022; Basan et al. 2020; Schink et al. 2022), which should result in an increase in R/F at the community level with increasing temperatures. If our hypothesis is correct, we should be able to predict changes in community composition across a temperature gradient.

## Methods

### Microbial communities enriched in defined media with different carbon sources across a temperature gradient

To get a diverse community of microbes as a source, we collected ∼1.5 g leaf litter from White Oak (*Quercus alba*) trees at Yale West Campus (41°15’27’’N, 72°59’34’’). The temperature at the time of sampling was 25°C. To create the inoculum, we added phosphate-buffered saline (PBS) to reach a final volume of ∼15 mL; we incubated the leaf sample for two days at room temperature vortexing the sample every day for 1 minute release the cells from the leaf. We let the leaves settle at the bottom and used the top liquid to inoculate 96 wells of five different deep-well plates (one per temperature). In each plate, we had wells with 500 µL of M9 minimal medium with one of 23 carbon sources (see Table S1). Each carbon source had the same concentration of carbon (0.07 C-mol/L). For each carbon source, we had four replicate wells. In addition, we had 4 wells with M9 and water as the control. In this paper, we are only examining the communities assembled in sugars (Table S1).

In the first two transfers, we added a final concentration of 200µg/ml of cycloheximide to exclude eukaryotic cells from our experiment (as in (Goldford et al. 2018)). We incubated each plate at a different temperature (12^°^C, 22°C, 30°C, 37°C, and 42°C) in different incubators. To control for variation in incubator conditions, we maintained our cultures at high relative humidity in big plastic containers inside each incubator. We tracked humidity and temperature conditions throughout the experiment with Track-It RH/Temp data loggers (Monarch instrument, NH, USA; Figure S1). We checked periodically to make sure that conditions were being maintained. The first 3 days, the 37°C treatment had consistently higher temperatures, between 38-39°C so we adjusted the settings in the incubator. Afterwards, it maintained a mean temperature of 37.7°C and a standard deviation of 0.15°C (excluding door openings). We kept the plates with new media at the appropriate temperature for at least one hour before the transfer to avoid large fluctuations in temperature.

After 48 hours of growth, we transferred 4 µL of each culture into 496 µL of fresh M9 media with the corresponding carbon source. We performed a total of 18 transfers to let the communities stabilize. We measured the optical density at a wavelength of 620nm (OD620) of each culture by sampling 100 µL cultures after each 48-hour growth cycle, and we stored the remaining cultures with a final concentration of 40% glycerol at −80°C. In previous work, we have seen that at 30°C twelve transfers are enough to allow communities to assemble into a generational equilibrium, where the community composition at the end of the batch incubation is maintained in subsequent incubations (Goldford et al. 2018). In this case, we allowed for 50% more passages to account for differences in growth rates across temperatures.

### DNA extraction and 16S rRNA gene sequencing

We extracted DNA from communities stored in 40% glycerol at −80°C after the last transfer. We let the plates completely thaw to remove the glycerol; we centrifuged the plates to pellet all the cells, and carefully removed all the supernatant. Then we re-suspended the cells in 180μl of enzymatic lysis buffer (20 mM Tris-HCl, 2mM sodium EDTA, 1.2% Triton X-100), we added 20 mg/mL of lysozyme from chicken egg white (Sigma-Aldrich) and incubated the mix at 37 °C for 30 min with gentle shaking (200rpm). To lyse the cells, we added 5μl of proteinase K and 200μl of buffer AL (Qiagen). We mixed the samples and incubated them for 1 hr at 56 °C. Following cell lysis, we extracted the DNA of all of these samples using the DNEasy Blood and Tissue kit (Qiagen) as described in the protocol. We stored the extracted DNA at −20 °C before shipping it in dry ice for sequencing. For sequencing, the 16S rRNA amplicon library was prepared as described in the Earth Microbiome Project Protocol (http://www.earthmicrobiome.org/protocols-and-standards/16s/). Briefly, the PCR-amplified libraries were prepared using the primer pair F515/R806. Then, the generated amplicons (spanning the V4 region of the 16S rRNA gene) were pooled and sequenced in paired-end reads using the Illumina MiSeq platform (Caporaso et al. 2012). All sequencing procedures including the library preparation were performed at the Argonne National Laboratory (Lemont, IL). Short-read sequences are available in the NCBI SRA, accession number (PRJNA992630).

### 16S sequencing data analysis

The first step to analyzing our 16S amplicon data was to demultiplex the file using *idemp* a custom script by Yinghua Wu (https://github.com/yhwu/idemp). We then used the pipeline from dada2 version 3.16 (Callahan et al. 2016). Briefly, we first filtered and trimmed reads according to their overall quality profiles. We trimmed the reads after 230bp for both forward and reverse reads or, if the quality was too low, after the first instance of a quality score less or equal to 11. After trimming forward reads with more than 1 “expected error” (EE) and reverse reads with more than 2 were discarded. DADA2 calculates expected errors based on the quality score Q, as *EE* = Σ(10^-Q/10^). Next, reads were de-replicated and merged. At this point reads should be ∼250bp, we removed reads longer than 256bp (0.17%). There were no reads shorter than 248. We also removed chimeras. After the whole process, we ended up with a median of 50,000 reads per sample (Figure S2). We assign taxonomy using the Silva database (version 132) using the RDP classifier (Wang et al. 2007) as implemented in DADA2. For all downstream analyses, we rarefied our data at the minimum number of reads per sample (8064) using the R package vegan (Oksanen et al. 2007).

### Respirator to fermenter ratio estimation

To estimate the R/F (respirator to fermenter) ratio from 16S data, we assigned members of the Enterobacteriaceae, Enterococcaceae, and Lachnospiraceae families to fermenters, and Pseudomonadaceae, Burkholderiaceae, Rhizobiaceae, Moraxellaceae, and Xanthomonadaceae to respirators according to the literature (Table S2). In previous work, we have confirmed the functional role of Enterobacteriaceae and Pseudomonadaceae, the two dominant families in our communities (Estrela et al. 2022). In addition, all of the fermenter isolates from this work (family Enterobacteriaceae) produced organic acids when grown in glucose confirming their fermentation ability.

### Isolation of strains from enrichment communities

To evaluate the role of thermal responses of individual isolates in predicting community assembly, we isolated strains from microbial communities stabilized in media with one of four sugars (glucose, fructose, galactose, ribose) at five different temperatures. We thawed the microbial communities from the last transfer stored at −80°C. After making sure that these communities were well mixed, we took a small sample diluted and plated it in chromogenic agar (Universal Chromoselect, Sigma Aldrich). Chromogenic agar allowed us to distinguish our fermenter taxa (blue, purple, and pink colors) from groups of respirators (white colonies). We selected two different colonies of each group per carbon source and temperature. In some cases, we were not able to isolate respirators because there were none on agar plates. We streaked each isolate and picked a single colony each time to remove possible contamination, we repeated this streaking at least two times per isolate. We inoculated a single colony of each of these isolates in a different well of a 96-deep-well plate with 500 µL of trypticase soy broth (TSB) media on each well. We let these isolates grow for 24hrs at 30°C. We stored the isolates in 40% glycerol at −80°C for further analyses. We confirmed our functional assignment by identifying each isolate through Sanger sequencing. Before all phenotypic measures, we plated all isolates to check for contamination and confirm their identity.

To evaluate how well our isolates represented the communities, we aligned the Sanger sequences of our isolates to amplicon sequencing reads (ESVs) of the communities as described in (Estrela et al. 2022). We performed a pairwise alignment between Sanger sequences and all ESVs from the same community, the alignment with the highest alignment score was picked with zero or one base pair mismatch. In general, our isolates represented their source communities well with an average coverage of 84.4% for replicate community 2 and 81.2%for replicate community 4 (Figure S3).

### Measurement of growth curves of strains

We performed growth assays with isolates under 12°C, 22°C, 30°C, 37°C, and 42°C. To acclimate the strains before growth assays, we grew isolates from glycerol stocks in 500 µL trypticase soy broth (rich media) in 96 deep-well plates for 24 hours in incubators with desired temperatures. We transferred 5 µL of these cultures to 200 µL fresh M9 media with one of the four carbon sources (glucose, galactose, fructose, and ribose) in 96-well NUNC plates. Medium with each carbon source had three replicates for experiments performed at 12°C, 22°C, and 30°C. One plate was made for each carbon source at 37°C and 42°C. After incubating 12 or 4 plates for 48 hours under desired temperatures, we measured OD values for grown cultures. Then, we transferred 4 µL of grown cultures to 200 µL fresh media. We used a plate reader (Biotek Epoch 2) with a microplate stacker (BioStack 4) to measure the growth curve for each isolate over the course of 48 hours in a temperature-controlled room set at 12°C, 22°C, or 30°C. The highest temperature for the temperature-controlled room was 30°C, so we measured the growth curve of isolates in four carbon sources in four plate readers set at 37°C or 42°C (accuSkan FC). To make sure there were no significant effects of plate reader in our growth measurements we compared data from both plate readers for plates at 30°C (Figure S4).

To determine growth parameters (lag phase, maximum growth rate) we used a custom function that fits a generalized additive model (gam) to the ln(OD620) value over time using a penalized cubic regression spline (bs=cs) using the gam function from the mgcv package (Wood 2011) in R version 4.2.2 (R Core Team 2023). These are very flexible models that allow us to track different moments in these curves accurately. The cubic spline penalty minimizes problems with noise and overfitting while still closely following the dynamics of otical density over time. Our function then approximates the derivative for the predicted values at each point using a finite difference interval (1 ∗ 10^!&^). The maximum growth rate is calculated as the maximum derivative after filtering initial values (first two hours) because at this point noise is large in proportion to the signal leading to spurious values of maximum growth rate.

To fit thermal performance curves, we used a Gaussian response to temperature, where:

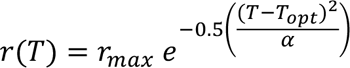

describes growth rate at a given temperature (*r(T)*) as a function of the maximum growth rate at the optimum temperature (T_opt_), the current temperature (T) in reference to the optimum, and a value of thermal breath (α) (Lynch and Gabriel 1987; Amarasekare 2015). From our fit we estimated the values of T_opt_, rmax, and α. We used the rTPC pipeline in R to fit this model to our data (Padfield, O’Sullivan, and Pawar 2021).

### Measurement of organic acid secretion by fermenters

We streaked isolates stored in 40% glycerol on chromogenic agar plates to check for contamination. After incubating the plates for one day in a 30°C incubator, we picked a single colony for each isolate and resuspended it in 200 µL PBS in a 96-deep-well plate. Then, we transferred 5 µL of these inocula into 500 µL fresh M9 media with glucose in five 96 deep-well plates. We incubated each plate under one of the five temperatures (12°C, 22°C, 30°C, 37°C, and 42°C). After two days, we transferred 5 µL cultures from each deep-well plate to 500 µL fresh M9 media with glucose in two replicate 96-well NUNC plates for destructive sampling at two time points. For temperature 12°C, we sample after 24 hours (T1) and 48 hours (T2). For the other four temperatures, we sacrifice one replicate plate after 12 hours (T1) and the other after 24 hours (T2). T1 is within the exponential phase of most cultures and T2 is a time when, on average, the population has reached carrying capacity in glucose, but bacteria have not switched to start consuming organic acids (Figure S5) (Estrela et al. 2022).

At each time point, we sampled 100 µL of culture to measure OD (620 nm). We centrifuged the rest of each culture at 3600 rpm for 25 minutes to separate cells and their supernatant. After centrifugation, we carefully transferred 200 µL of supernatant into 96-well plate filters and collected the supernatant into 96-well NUNC plates by centrifuging at 3200 rpm for 10 minutes. We stored the supernatant plates at −20°C until analysis. To obtain per capita organic acid production, and estimate yield, at T2, we streaked cultures from selected wells with fermenters on chromogenic agar plates for colony counting. We estimated all fermenter cell numbers at T2 by measuring OD (620 nm) values and the linear regression between colony numbers and OD (620 nm) values of selected wells (R^2^=0.939).

We used colorimetric methods to measure acetate and lactate concentrations in wells with fermenters based on the manufacturer protocols (abcam ab204719 for acetate, abcam ab83429 for lactate). We diluted samples to the range of the standard curve. We made two standard curves with identical standard dilutions for each batch of samples to cover variations among kits. We calculated final concentrations based on the sample dilution rate and the standard curve, a linear regression between measured OD (450 nm) and acetate amount of the corresponding standard.

### Measurement of glucose consumption

We measured the glucose consumption of all isolates across conditions using a Glucose (GO) Assay Kit (Sigma-Aldrich). First, we obtained 200µL of the supernatant at the endpoint by centrifuging at 3200 rpm for 10 mins using filter plates (Pall Acroprep, supor filter plate 0.25μm). Afterward, we prepared a glucose standard curve as specified by the assay kit (with a working range of 20 - 80 ug/mL), and we diluted the supernatants accordingly. Using a 96-well pipettor to minimize variation in reaction time, we added 20 uL of the dilutions and standards to another 96-well plate containing 40 uL of a colorimetric assay reagent. We incubated the plate at 37°C for 30 min. We carried out all the steps in complete darkness. We finally interrupted the reaction by adding 40 uL of 6N H2SO4 and measured the absorbance at 540 nm (Multiskan FC, Thermo Fisher Scientific). We calculated the absolute glucose consumption, as well as the percentage of glucose consumption based on the M9 medium (0.21% Glucose).

### Calculation of glucose efficiency

In previous work we proposed a model to describe the dynamics of these communities and quantitatively predict the ratio of respirators to fermenters, R/F (Estrela et al. 2022). Assuming that glucose is only consumed by fermenters (F) and the organic acids are only consumed by the respirators (R), we can approximate R/F as:

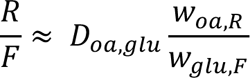

Where *W_glu,F_* is the yield efficiency of fermenters in glucose, *W_oa,R_* is the yield efficiency of respirators in organic acids, and *D_oa,glu_* is the number of acetate molecules that fermenters secrete back into the environment per molecule of glucose consumed. Thus, assuming that all carbon uptaken by *F* is either converted to biomass or secreted as organic acids, *D_oa,glu_* gives an idea of *F* carbon efficiency. A high value of *D_oa,glu_* means that most of the glucose uptake by *F* is secreted back to the media and therefore is no longer available for *F* biomass.

These assumptions are justified by a growth advantage of *F* over *R* in sugars and of *R* over *F* in organic acids, as well as the observation that *F* does not co-utilize sugars and acetate, instead switching between these carbon sources requires a shift from a glycolytic direction of central metabolism to a gluconeogenic direction leading to long lag times (Basan et al. 2020; Schink et al. 2022). In addition, in this work, we assume that organic acids comprise only lactate and acetate. These are the dominant carbon sources present in our communities (Estrela et al. 2022) and bacteria with a glycolytic preference (F) take a long time to switch and consume both of these carbon sources (Basan et al. 2020).

We obtained the empirical values of glucose efficiency (*D_oa,glu_*) at all experimental temperatures by dividing the total carbon consumed as glucose by our fermenter isolates over the total carbon secreted as organic acids by the same fermenters. Glucose efficiency is then 1-*Doa,glu*.

### R/F prediction

Using these empirical values of *D_oa,glu_* and our model, we can predict R/F. We can calculate empirically the biomass of fermenter taxa produced per carbon in glucose (*W_glu,F_*), as well as the amount of respirators biomass produced per carbon available as organic acids (*W_oa,R_*). To obtain *W_glu,F_*, we obtained the optical density after cultures reached their carrying capacity (the same time at which we measured glucose and organic acids in the supernatant). We removed the blank from those values and then divided it by the amount of carbon consumed in the form of glucose. To avoid artifacts of isolates growing at temperatures different than their original community, we used only the values of isolates growing at the temperature at which they were isolated.

We determined *W_oa,R_* from the optical density of communities assembled in acetate (0.07M of carbon). These communities were composed in their majority by respirator taxa (median respirator relative abundance: 0.97, Q1: 0.75, Q3: 0.99), so we could multiply the optical density by the relative abundance of respirators to get an estimate of overall biomass. Then, we calculated the difference between final and initial densities, and assumed that all of the acetate had been consumed. Thus, we calculated *W_oa,R_* as the difference between the final and initial optical densities (the initial optical density is the final optical density divided by dilution factor of 125) divided by the total amount of carbon present in the media as acetate. That is: *W_oa,R_* = (*OD_final_* − *OD_initial_*)/*C_acetate_*.

Finally, as described in our mode (Estrela et al., 2022), we obtained the predicted values of R/F by dividing the values of *D*_1(,345_ for each isolate, by the *W*_345,7_ calculated for the same isolate. We then multiplied all of these values by the four possible values of *W*_1(,6_ (one for each replicated community).

## Results

### Community composition changes with temperature in a predictable manner

We collected a diverse microbial community from decomposing leaf litter and used it to inoculate replicate habitats with one out of 16 sugars as the only supplied carbon source. To evaluate how temperature affects community assembly in different sugars, after inoculation, we let the communities stabilize through serial transfers in each carbon source at each of five different temperatures (12.5°C, 22°C, 30°C, 37.7°C, 42°C) (Figure 1A; Methods).

**Figure 1.**
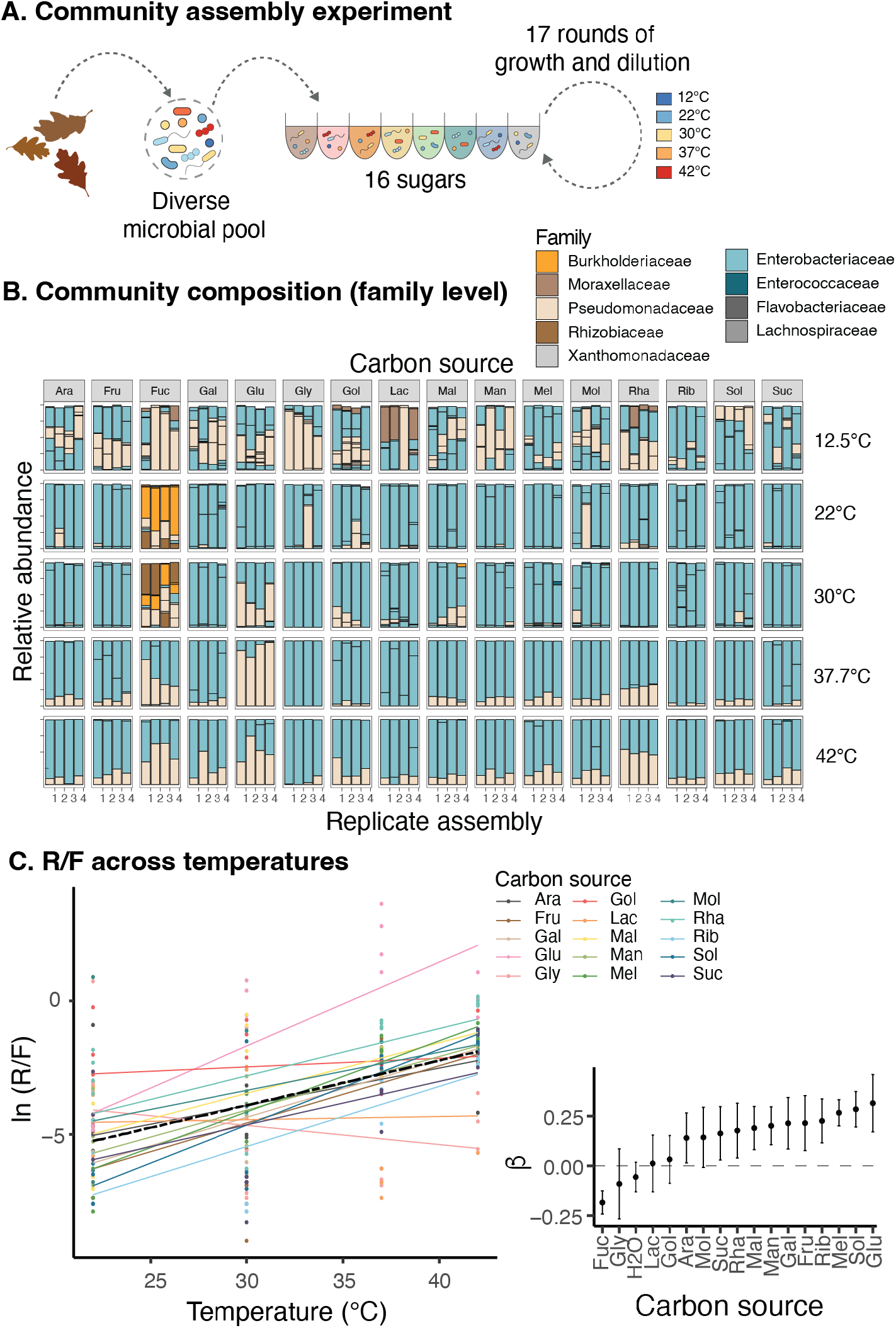
Community assembly experiment. A. Schematic of the protocol. We inoculated different carbon sources with a diverse pool of microbes and let them grow at one of 5 temperatures. B. Relative abundances of different families in each of the communities. The columns are the different sugars and the rows different temperatures of assembly. The 4 bars within the same row and column are replicated assemblies. C. Relationship between temperature (from 22°C to 42°C) and ln(R/F) across the 15 sugars. The dashed line is the mean relationship across carbon sources. Fucose was excluded from this analysis because it had very distinct dynamics than all of the other sugars. C. Estimates of the slopes (and 95% CI) for the relation between temperature (from 22°C to 42°C) and ln(R/F). Fucose and water are included. We did not observe a significant relationship between temperature and ln(R/F) in communities assembled in water without supplemental carbon sources.

Within each carbon source and temperature the replicate communities were fairly consistent at the family level (Figure 1B). Moreover, we saw replicable patterns across temperatures for all communities. These communities assembled in different sugars were dominated by fermenter taxa (members of the Enterobacteriaceae family) followed by respirators of Pseudomonadaceae, consistent with previous observations (Figure 1B) (Goldford et al. 2018; Estrela et al. 2022), and, as in previous work, despite replicability at the family level, community assembly showed little convergence at the level of exact sequence variants (ESVs) (pairwise Bray-Curtis distance between replicates: Q1 = 0.097, Median = 0.371, Q3 = 0.53; pairwise Bray-Curtis distance between sugars: Q1 = 0.276, Median = 0.619, Q3 = 0.896). This divergence between replicates is highest at 12.5°C and decreases with increasing temperature (Figure 1B, Figure S6).

Consistent with our hypotheses, excluding fucose, the respirator to fermenter ratio (R/F) showed an increase with increasing temperatures from 22°C to 42°C (Figure 1C). After log transforming the data, we observed a slight positive relationship between temperature and ln(R/F) (methods; β_1_ = 0.17 ± 0.016 SE, p-value < 0.001; Figure 1C). This relationship is maintained for most of the sugars evaluated (Figure 1C, D). Glycerol, galacticol, lactose, arabinose, and sucrose show no clear relationship between temperature and R/F (95% confidence interval of the slope includes 0). Only fucose showed a significant negative relationship between temperature and ln(R/F) (β = −0.184, 95%CI [−2.42, −0.13]). In this sugar we also observed lower optical densities (Figure S7), and overall a much more distinct pattern of assembly (even at the family level) (Figure 1B, D).

One possibility that could explain the overall increase in respirators with increased temperatures (after 22°C) is if, in our original inoculum, respirative bacteria had a fitness advantage over fermentors as temperature increases. However, across all temperatures, fermenters tended to grow faster than respirators in the four sugars tested (glucose, fructose, galactose, ribose) (Figure 2, Figure 3A). After controlling for the temperature of isolation and carbon source, the difference in growth fermenters and respirators in the supplied sugars increased approximately 1.6 times between 22°C and 30°C (<u>r</u>(22)_F_ − <u>r</u>(22)_R_ = 0.082 ± 0.005*SE* and <u>r</u>(30)_F_ − <u>r</u>(30)_R_ = 0.131 ± 0.005*SE*; Figure 2), and the R/F ratio for these same carbon sources increased from a mean of 0.012 at 22°C to 0.3 at 30°C.

**Figure 2.**
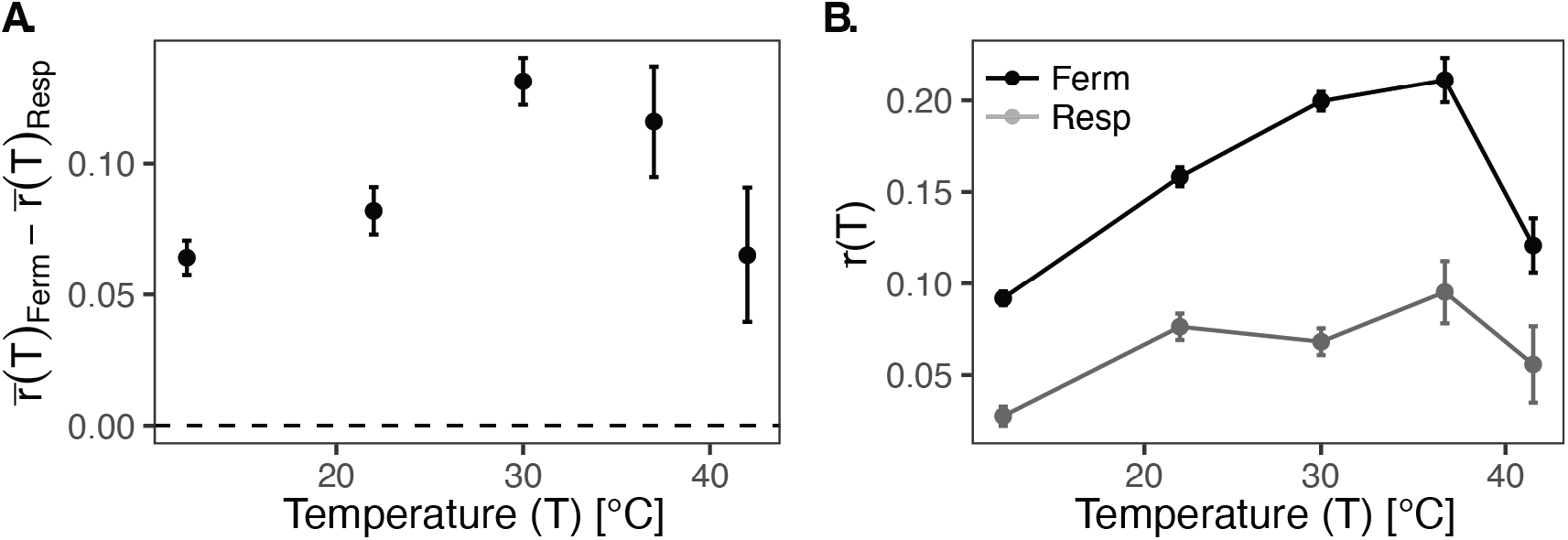
Estimated marginal mean growth rate (<u>r</u>(T)) after accounting for the temperature of community assembly from which each isolate was taken and the carbon source in which growth was measured. **A.** Difference between the estimated mean growth rate of fermenters and respirators at each temperature. **B.** The estimated mean growth rate of fermenters (ferm) and respirators (resp) at different temperatures. Error bars are the 95% confidence intervals for each estimate.

**Figure 3.**
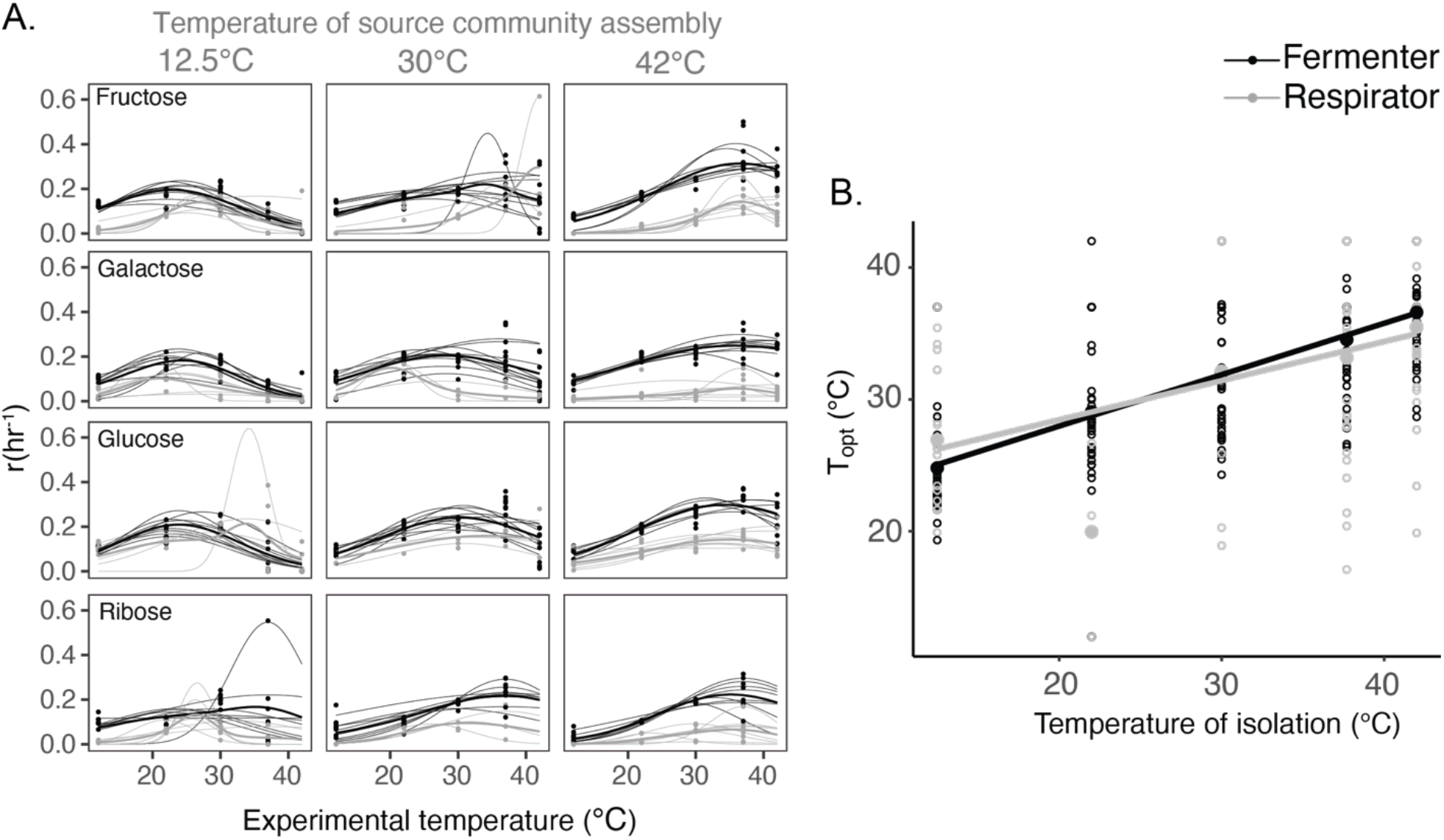
Response of growth rates of fermenters and respirators to increasing temperatures. A. Each column corresponds to a source temperature (the temperature at which the communities were assembled, only 12.5, 30, 42 are shown, all plots are in supplementary), and each row corresponds to the sugar used to measure growth. Each thin line is the Gaussian model fit to one isolate, thick lines are the mean for fermenters (black) and respirators (gray). B. Relationship between the temperature of isolation (the temperature at which the source community was assembled) and the optimal temperature of that isolate (T_opt_). Outlined small circles are the values of each isolate and filled big circles are the mean value. Lines are the slopes from a linear model.

In contrast, at 12.5°C we observed high diversity, lower replicability between replicates (pairwise Bray-Curtis distance between replicates: Q1 = 0.46, median = 0.501, Q3 = 0.54; Figure S5), and a high R/F ratio across all carbon sources (median = 0.75, Q1 = 0.32, Q3 = 1.47, N = 64). At this temperature the difference in growth between respirators and fermenters is small (r(T)_F_ − r(T)_R_ = 0.064, 95%CI [0.058, 0.071]; Figure 2A). As temperatures increase the difference in growth between respirators and fermenters increases to a maximum around 30°C and decreases again afterward, but even at 37.7°C the growth difference between functional groups is almost twice the 12°C value (r(T)_F_ − r(T)_R_ = 0.116, 95%CI [0.095, 0.137]; Figure 2A). This difference in growth between fermenters and respirators across temperatures is mostly shaped by changes in growth rates of fermenters across this same temperature gradient (Figure 2B). Respirators growth is low across all temperatures even though it still increases slightly until 37°C and then goes down again at 42°C (Figure 2B).

### Community assembly at higher temperatures selects isolates with higher T_opt_

We observed differences in maximum growth rates across functional groups (fermenters have higher maximum growth rates across carbon sources), as well as differences in maximum growth rate across carbon sources, and the temperature of origin. We also observed a significant interaction between functional group and temperature of origin on maximum growth rates (Table S3; Figure 3A, Figure S7).

The process of community assembly sorted isolates of the same functional groups according to their thermal physiology. That is, strains isolated from communities assembled at lower temperatures tended to have a lower thermal optimum (T_opt_) than those from communities assembled at higher temperatures (Figure 3). For each degree higher in the temperature at which communities were assembled, fermenter isolates had on average a 1.073°C±0.107SE increase in T_opt_ (with an intercept of −5.12°C; Figure 3B). Respirators had a similar slope (0.756°C±0.123SE; p-value > 0.05) but a larger intercept (Ferm-Resp: 12.626°C±0.123SE, p-value = 0.018; adjusted R^2^ = 0.35; Figure 3B).

### Fermenters face a trade-off between growth and glucose efficiency at higher temperatures

We hypothesized that the increase in R/F with temperature is the result of a decrease in metabolic efficiency with faster growth rates (Figure 4A). Fermenters from communities previously assembled in glucose at 30°C showed increased organic acid production at 28 hrs with increased growth rates (slope: 21.67±3.559SE mM secreted to media per hour increase in growth rate; R2 = 0.44, p-value < 0.0001) whereas respirators do not secrete acetate at any growth rate (slope:-0.31±0.45SE mM secreted to media per hour increase in growth rate; R2 = 0.03, p-value = 0.5); Figure 4B). Therefore, we expected that increasing growth rates with increasing temperatures would lead to greater amounts of acetate secreted (Figure 4A), but we did not know if different temperatures would have different slopes.

**Fig 4.**
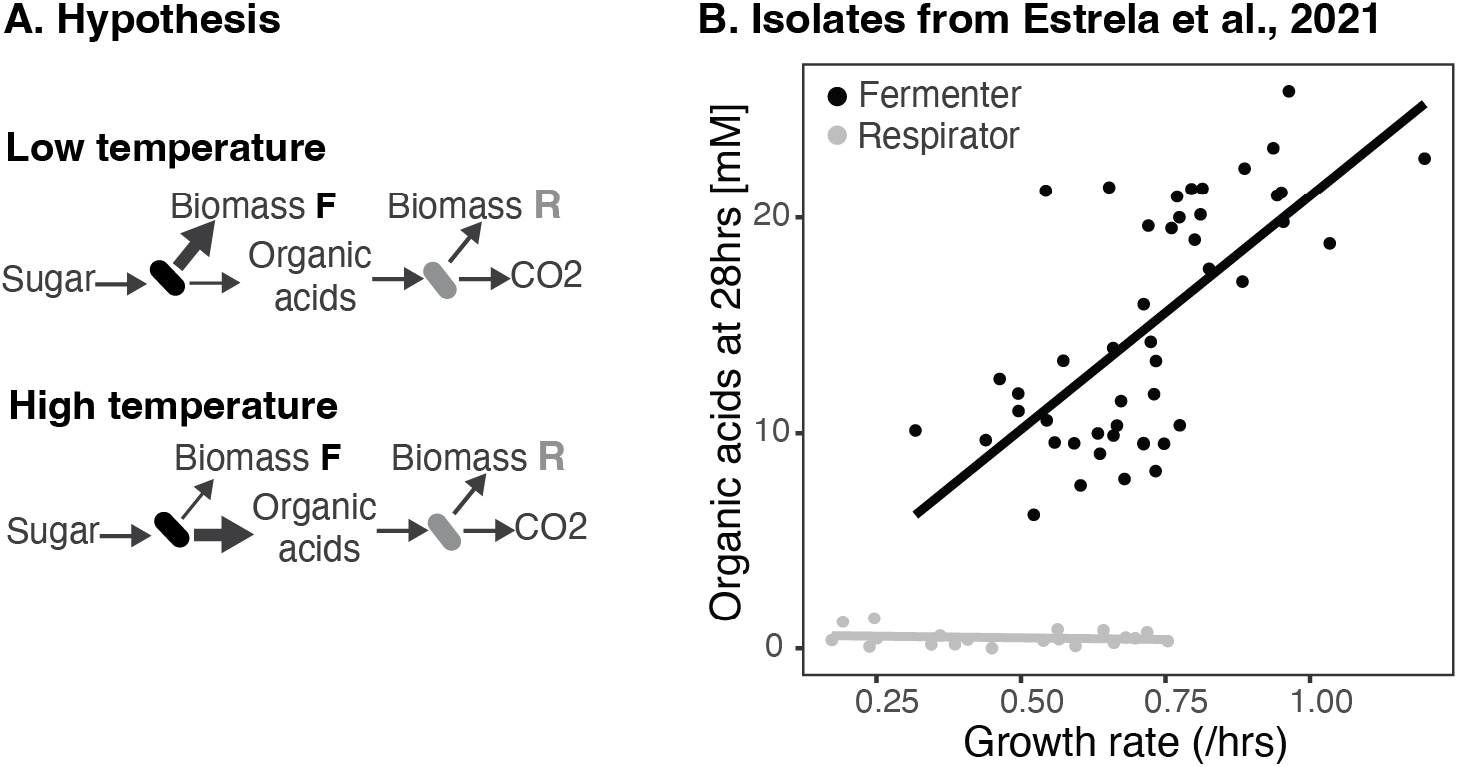
In our conceptual model of interactions between functional groups, fermenters (F) consume sugars and use them for biomass production or secrete them as organic acid waste products into the media. Respirators (R) consume these organic acids and make biomass or produce CO_2_. As temperatures increase we expect that fermenters will release more organic acids into the media as a result of their increased growth rate (A). This hypothesis is based on previous data of isolates from communities assembled of glucose at 30°C. Fermenters from these communities secrete an increased concentration of organic acids into the media with increasing growth rates (B).

We grew all isolates in minimal media with glucose and measured the concentration of organic acids (acetate and lactate) produced by the time fermenters reach their carrying capacity (Figure S4). We observed a linear relationship between growth rate and the secretion of organic acids (Figure 5A). For each unit of density (ln(OD600)) per hour faster in growth, we observed an increased total concentration of carbon secreted as organic acids of 68.98± 7.01 SE mM (Fig5A;p <0.0001; adjusted R^2^ = 0.31). This model, however, is obscured by highly variable (and often high) secretions after the T_opt_. If we remove all points above the T_opt_, we get carbon secreted as organic acids per hour faster in growth of 96.49± 7.5 SE mM (Figure 5A). The total amount of carbon secreted as organic acids increased with temperature (Figure 5B), reaching a maximum at 37°C and then decreasing at 42°C (after Topt for most isolates). Overall, after 37°C (when most isolates have passed their T_opt_) secretions stay high for the drastic decrease in growth (Figure 2B; Figure 5A,B).

**Figure 5.**
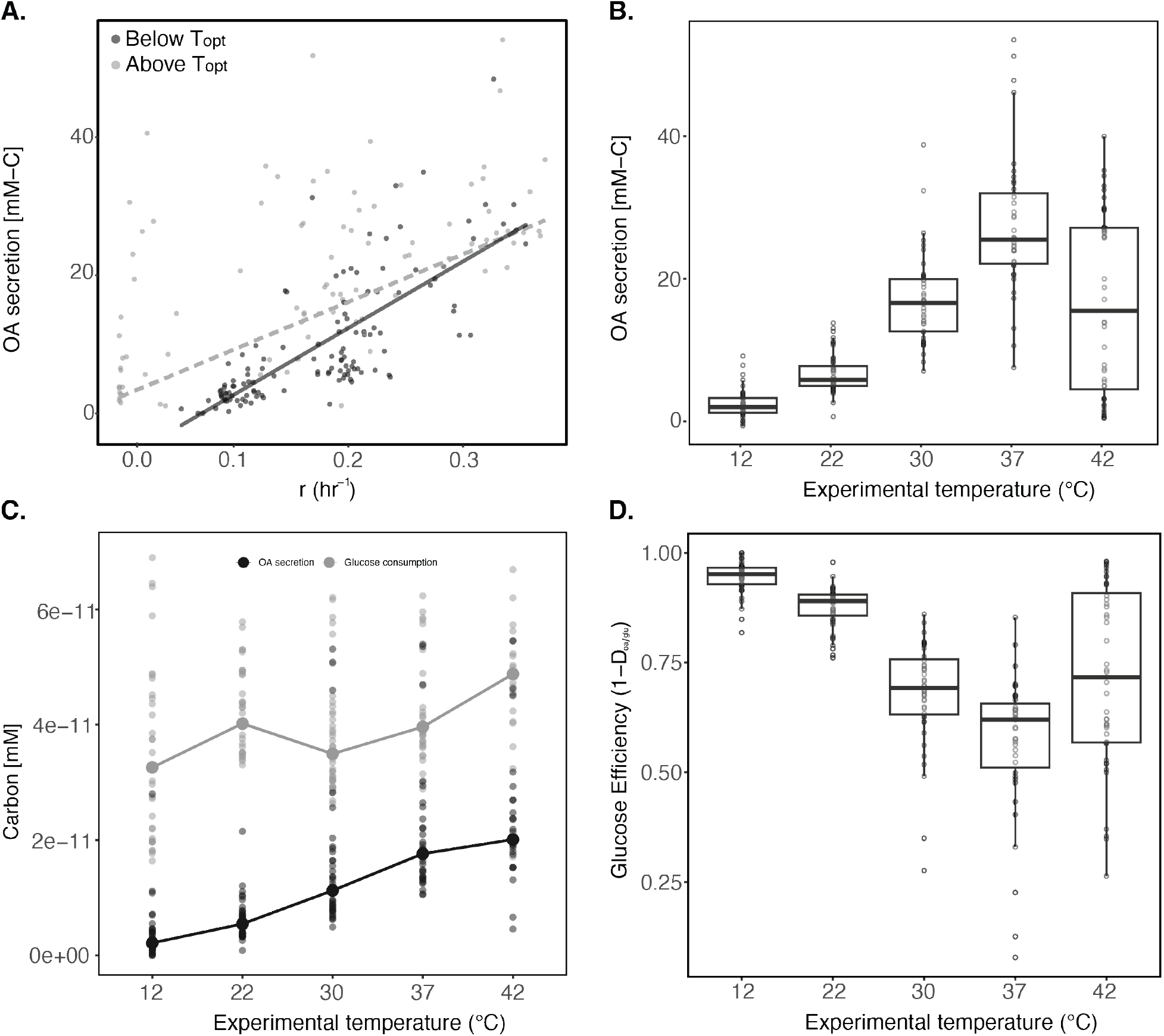
Effect of temperature on the carbon use of fermenters after glucose depletion. **A.** Total carbon that is secreted as organic acids (OA) at different growth rates, with points colored by their experimental temperature above (light) or below (dark) the thermal optimum. Trendlines show linear fits to all data (dashed) and data below the thermal optimum (solid). **B.** Total secreted carbon per temperature. **C.** Consumed and secreted carbon per cell at different temperatures. **D.** Glucose efficiency, calculated as 1 - D oa/glu (ratio of secretions per unit of glucose consumed by fermenter cells), at different temperatures.

The R/F ratio can be quantitatively predicted from a simple model describing the biomass production of fermenters based on the conversion of glucose into biomass by fermenter taxa, and biomass production of respirators from consuming organic acids secreted by fermenters (Figure 4A, Estrela *et al*., 2021). We wondered if this simple model could also predict the changes in the R/F ratio across a temperature gradient. According to this model carbon from glucose can be utilized for biomass formation of fermenters or released into the media as organic acids. Thus, the fraction of organic acids produced per molecule of glucose (*D*_oa,glu_) is related to the amount of carbon used in biomass (1-*D_oa,glu_*) or glucose use efficiency.

We observed a trade-off between growth rate and glucose efficiency with increasing temperatures leading to faster growth rates at the cost of secreting more organic acids. From glucose consumption and organic acid secretions, we calculated the glucose efficiency (1-*D_oa,glu_*; see methods). Glucose efficiency decreases with increased temperature and then increases slightly at 42°C (Figure 5D). Glucose efficiency is shaped by secretion of organic acids and a significant increase of glucose consumption per cell at 42°C. While the secretions per cell increase steadily with temperature linear model with temperature of isolation as random factor:5.08 *10^-12^± 1.22*10^-12^ mM/(cell*°C), p-value<0.0001; Figure 5C), glucose consumption shows a less clear trend that disappears when removing the 42°C data (linear model with temperature of isolation as random factor: 5.17 *10^-11^± 1.58*10^-11^ mM/(cell*°C); p-value=0.0012; removing 42°C data: 1.54 *10^-12^± 2.311*10^-12^ mM/(cell*°C); p-value=0.5).

From 12°C to 37°C, as temperatures increase, we see increased growth rates, a maintenance of glucose consumption (Figure 5C), and an increase in organic secretions per capita. All consistent with a decrease of glucose use efficiency (Figure 5D). In contrast, the seemingly increased in glucose efficiency at 42°C is misleading: there is more glucose consumed per molecule of organic acid secreted, but temperature-induced mortality, also means that there is less biomass sustained. At 42°C most isolates are past their T_opt_ and therefore, for the same growth rate they show a much higher secretion of organic acids (Figure 5A; Figure S8). Altogether, these results are consistent with the observed linear increase in R/F ratio with temperature in glucose.

### Changes in the fermenter’s glucose efficiency predict changes in R/F across temperatures

According to our model (see methods; Estrela et al., 2021), R/F can be predicted from three parameters: the biomass of respirators produced per carbon consumed in organic acids (*w_oa,R_*), the biomass of fermenters that is produced per carbon in glucose consumed (*w_glu,F_*), and the amount of carbon as organic acids that is secreted by carbon of glucose consumed (*D_oa,glu_*). We obtained empirical measurements of *w_glu,F_* and *D_oa,glu_* from our physiological measurements with fermenters in glucose, and in the absence of empirical measurements of respirators growing in organic acids, we obtained *w_oa,R_* from the optical density and relative abundance of respirators in communities assembled in acetate at different temperatures.

Our predicted R/F does not match our observed values at 12°C, given the overall low biomass of respirators and very low *D_oa,glu_* we predicted a R/F value of 0.0015 (Q1:0.0009, Q3:0.0024), but overall our communities have a much higher R/F of 0.698 (Q1: 0.299, Q3:1.328). After 12°C we predicted an increase of R/F with temperature until 37°C (Median R/F: 0.714, Q1: 0.707, Q3: 0.903) and then it should be mostly maintained at 42°C (Median R/F: 0.776, Q1: 0.647, Q3:0.828). This is consistent with the behavior of R/F in glucose communities (the sugar we used for our measurements). Quantitatively, however, our predictions tended to overestimate R/F for other sugars, and in the case of glucose, our predicted ratio overestimated the relative abundance of respirators at 22°C and underestimated them at 37°C. Our model accurately predicted R/F of glucose communities at 30°C and 42°C (Figure 6). Using coarse measurements of respirator biomass and measuring all parameters from individual isolates (not in a community context) we obtained a prediction that qualitatively follows the dynamics of our communities from 22°C to 42°C, and quantitatively sits in between the observed values for all communities and the ones assembled in glucose.

**Fig 6.**
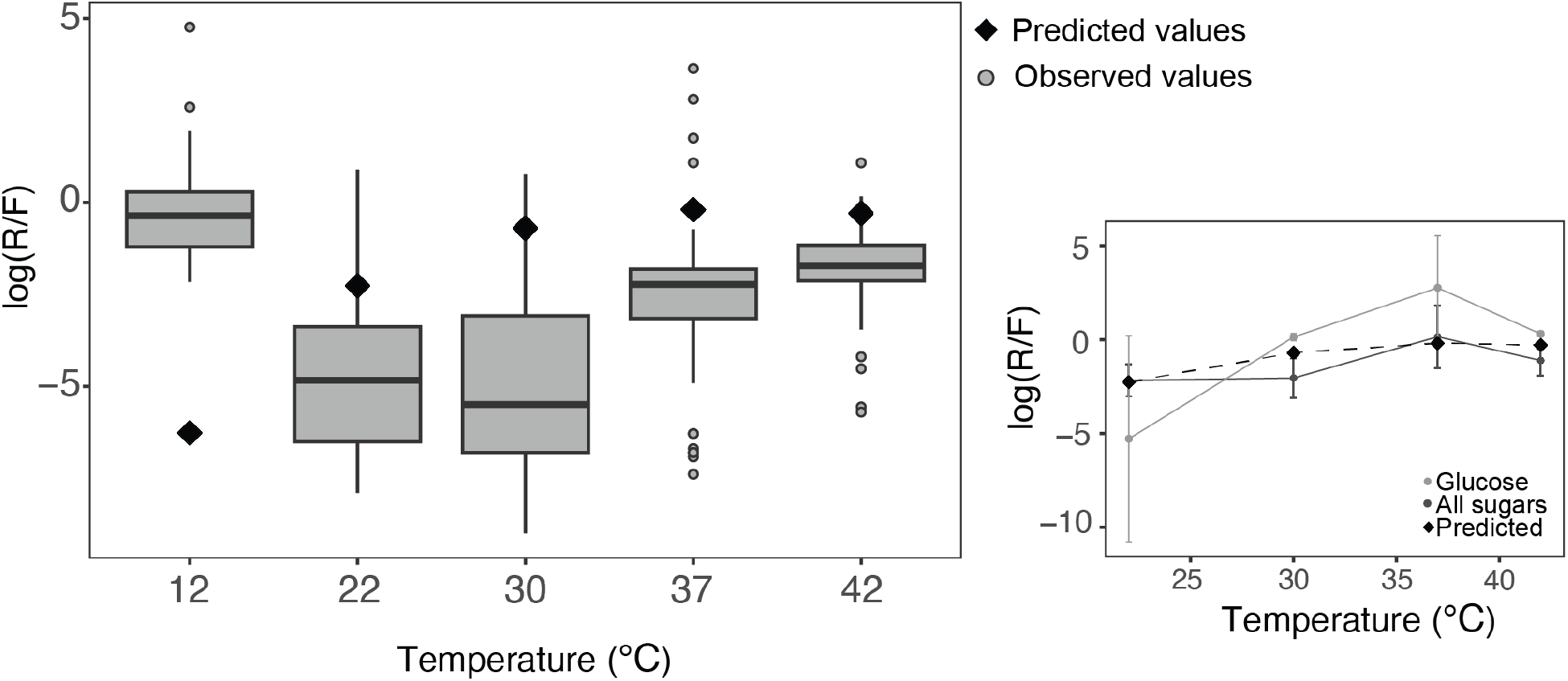
Effect of temperature on the carbon use of fermenters after glucose depletion. **A.** Total carbon secreted as organic acids (OA) at different growth rates, with points colored by their experimental temperature above (light) or below (dark) the thermal optimum. Trendlines show linear fits to data obtained at temperatures above the thermal optimum (dashed) and below (solid). **B.** Total secreted carbon per temperature. **C.** Consumed and secreted carbon per cell at different temperatures. **D.** Glucose efficiency, calculated as 1 - D oa/glu (ratio of secretions per unit of glucose consumed by fermenter cells), at different temperatures.

## Discussion and conclusions

Microbial communities often show high variation at lower taxonomic levels consistent with significant contributions of stochasticity (Albright, Chase, and Martiny 2019), priority effects (Debray et al. 2022; Chappell et al. 2022), and alternative stable states (Estrela et al. 2022). Nevertheless, these same communities are often remarkably replicable in function (Louca, Parfrey, and Doebeli 2016; Nelson, Martiny, and Martiny 2016; Burke et al. 2011).

In previous work, we have shown that we can derive simple rules of community assembly and quantitatively predict the relative abundance of different functional groups (fermenters, and respirators) using some basic knowledge about the physiology and metabolism of these groups. In this study, we were able to expand these simple rules to a temperature gradient.

Consistently with previous work (Basan et al. 2015), we have previously observed that fermenters display a trade-off between growth and yield, increasing their organic acid secretions with increasing growth rates (Estrela et al. 2022). Here, we showed that with increasing growth rates resulting from higher temperatures, there is an increase in organic acid secretions, and therefore a decrease in glucose use efficiency. Overall, this means proportionally more resources available for respirator taxa, and an increase in the ratio of respirators to fermenters (R/F).

With the exception of communities assembled at 12°C, using physiological data of fermenter isolates growing in glucose, and biomass data from communities assembled in acetate, we were able to predict qualitatively the patterns of change of R/F. Our quantitative predictions tended to overestimate R/F for most carbon sources but performed better for communities assembled in glucose (except the ones at 37°C) where communities in glucose had a really high R/F. Overestimation is to be expected given that our respirator biomass comes from communities starting at high concentrations of acetate. In contrast, in our sugar communities organic acids reach much lower concentrations of organic acids, and only after fermenters have depleted most of the glucose. Thus, and given that growth rate is a saturating function of resource concentration, it would be expected that respirators cannot achieve the same biomass per molecule of organic acid in the sugar communities.

All of the communities in this study were assembled from the same original pool of species, nevertheless, we observed distinct community composition at the ESV level, even between different replicate assemblies in the same carbon source and at the same temperature. This result indicates a significant contribution of stochasticity in community assembly. Despite stochasticity, we observed functional replicability across communities and a clear effect of selection on community assembly. For example, isolates with different thermal optima were favored at different temperatures.

The communities assembled at 12°C are the exception. These communities have high diversity within and between communities, even at high taxonomic levels. These patterns are consistent with expectations under a neutral regime (Thompson et al. 2020; Gravel et al. 2006; Vellend 2010) and at this temperature, the difference between functional groups is minimized. Fermenters grow only slightly faster than respirators and secrete little to no organic acids into the media (even after 48hrs of growth).

Our results demonstrate how physiological and metabolic constraints can be used to predict the effects of environmental factors on community assembly. In this case, two simple metabolic constraints help us predict changes in community assembly along a temperature gradient. First, fermenters grow faster in sugars than respirators, but once they are growing in sugars it takes them a long time to switch strategies and start consuming organic acids (Estrela et al. 2022). Second, increasing the fermenters’ growth rates leads to increasing organic acid secretions (Estrela et al. 2022; Basan et al. 2015).

The first of these constraints seems to result from the underlying metabolic architecture. Sugars and organic acids get catabolized by the central metabolism, using opposite directions of the same metabolic pathways. This metabolic structure restricts the co-utilization of organic acids and sugars leading to conflicts between pathways and potential futile cycles. This metabolic architecture imposes a trade-off between growth rates on one kind of carbon source and the time required to switch metabolic pathways (Schink et al. 2022; Basan et al. 2020), and could have favored the evolution of specialist taxa (e.g. fermenters vs. respirators). In particular, fermenter (glycolytic) taxa grow fast in sugars but have a long lag time between exhausting the sugar and before they can grow on available organic acids (Basan et al. 2020; Schink et al. 2022). Specialization along this axis seems to play an important role in structuring different microbial communities. In a recent study, for example, researchers found that most of the variation between metabolic preferences of isolates from marine heterotrophic bacterial communities was explained by an axis of sugars to organic acids preference (Gralka, Pollak, and Cordero 2022). In our system, this preference underpins a clear distinction between communities assembled in sugars or in organic acids (Estrela et al. 2021).

The second of these constraints (i.e. faster growth of fermenters is associated with increasing organic acid secretions) comes from previous observations of growth dependency of overflow metabolism (the production of seemingly wasteful by-products during growth on glycolytic substrates even in the presence of oxygen, when full respiration would be more efficient) (Basan et al. 2015; Millard et al. 2021; Szenk, Dill, and de Graff 2017). We had previously observed this relationship between growth rates and overflow metabolism across fermenter species in our system when grown at 30° but we did not know if this relationship would be maintained across temperatures.

It has been proposed that overflow metabolism increases with growth rates due to increasing proteome limitation (fermentation is less efficient in terms of energy conversion but it is cheaper in terms of proteome use) (Basan et al. 2015). In conflict with our results, these models predict that increased translational efficiency at higher temperatures compensates for the demands of faster growth, maintaining the same levels of overflow metabolism along a broad range of temperatures (Mairet, Gouzé, and de Jong 2021). Here we show an increase in organic acid secretion with increasing temperatures, and, except for isolates growing past their T_opt_, all temperatures fall along the same line.

There are multiple possible reasons for this discrepancy between theoretical predictions and our observations. First, there are other possible mechanisms predicting overflow metabolism, all based on growth constraints (de Groot et al. 2020). Second, while our environments cannot be fully oxygen limited or we would not see growth of respirators, our media is never shaken and therefore there is spatial and temporal variability in oxygen availability. In addition, changes in temperature affect rates of oxygen consumption and oxygen solubility, potentially affecting the metabolic strategies of fermenters. A better understanding of overflow metabolism requires studying different organisms in a range of environmental conditions. As such, not only can physiological studies provide insight into ecology, but a better understanding of microbial ecology and evolution can provide insight into bacterial physiology.

Finally, this study demonstrates the importance of considering metabolic plasticity, and different metabolic strategies when trying to understand the thermal responses of microbial communities. Bacteria are able to change their metabolic strategies in response to environmental conditions and changing constraints (Basan et al. 2015). For instance, microbes can decrease their investment in biomass to cope with increased maintenance costs under stress conditions (Malik et al. 2020). With increasing temperatures, cells will experience protein unfolding-related stress (Richter et al. 2010) which (at very high temperatures) will stimulate cell allocation towards less biomass investment (Mairet, Gouzé, and de Jong 2021). Even before unfolding related stress sinks in, it has been hypothesized that at higher temperatures carbon use efficiency (CUE) should decrease due to higher thermal sensitivity of catabolism than growth (Allison 2014; Manzoni et al. 2012; Ye et al. 2020). However, the evidence is mixed at population and community levels (Frey et al. 2013; Ye et al. 2019; T. P. Smith et al. 2021). Our work suggests that part of the answer might be related to the expression of different metabolic strategies in response to increasing temperatures and changing carbon sources.

Consistent with these observations, long-term warming experiments suggest that changes in dominant carbon sources might have affected the carbon use efficiency of microbial communities in the warming treatment (Melillo et al. 2017). Similarly, a meta-analysis showed variation in CUE with carbon source additions, demonstrating a significant difference between organic acids (excluding amino acids) and glucose (Qiao et al. 2019). These studies highlight the need for systematic physiological descriptions and the development of theories that integrate information from cells into communities and ecosystem processes.

Long-term changes of microbial communities in response to climate change in natural ecosystems such as soil are more complex than our communities assembled in synthetic habitats. However, the relatively simple conditions in the lab make it possible to control and monitor community assembly under different conditions, resulting in a mechanistic understanding of their response to focal environmental factors (Estrela, Sánchez, and Rebolleda-Gómez 2021; Sun and Sanchez 2023). Ecological forecasting is a challenging task. Even the integration of more detailed physiological descriptions remains a challenge. First, we require a better understanding of how physiological changes at the organismal level can affect species interactions. Second, it would be intractable to incorporate detailed physiological information for all species and interactions. Instead, in this paper, we use broad functional responses and general trade-offs to increase our predictability.

These communities can potentially serve as representations of some simple environments, but their main role is to illuminate important processes potentially affecting community structure and function. These simple replicated communities represent a proof-of-principle that a better understanding of bacterial metabolism and physiology can provide simple rules to predict changes in community composition in response to abiotic changes.

## Data and Code Availability

All data and code are available on a repository on GitHub (https://github.com/mrebolleda/TemperatureCommunities).

