## Supplementary information for "Predictive microbial community changes across a temperature gradient"

**Table S1.** Carbon sources used in community assembly experiment.

| Carbon source | Starting compound | Carbon source class | Molecular mass | Carbon atoms per molecule |
|---|---|---|---|---|
| Glucose | Glucose | Sugar | 180.156 | 6 |
| Maltose | Maltose | Sugar | 360.31 | 12 |
| Lactose | Lactose | Sugar | 342.3 | 12 |
| Melibiose | Melibiose | Sugar | 342.3 | 12 |
| Galactose | Galactose | Sugar | 180.16 | 6 |
| Sucrose | Sucrose | Sugar | 342.3 | 12 |
| Fructose | Fructose | Sugar | 180.16 | 6 |
| Mannose | Mannose | Sugar | 180.16 | 6 |
| Ribose | Ribose | Sugar | 150.13 | 5 |
| L-Arabinose | L-Arabinose | Sugar | 150.1 | 5 |
| Mannitol | Mannitol | Sugar alcohol | 182.17 | 6 |
| Sorbitol | Sorbitol | Sugar alcohol | 182.17 | 6 |
| Galacticol | AKA Dulcitol | Sugar alcohol | 182.17 | 6 |
| Glycerol | 80% Solution | Sugar alcohol | 92.09 | 3 |
| Fucose | Fucose | Sugar (uncommon, part of some cell-surface glycans) | 164.16 | 6 |
| Rhamnose | Rhamnose | Sugar | 182.17 | 6 |
| Oxoglutarate | 2-ketoglutaric acid disodium salt dihydrated | Organic acid | 226.09 | 5 |
| Pyruvate | Sodium Pyruvate | Organic acid | 110.84 | 3 |
| Oxaloacetate | Oxaloacetate | Organic acid | 132.07 | 4 |
| Fumarate | Sodium hydrogen fumarate | Organic acid | 138.05 | 4 |
| Succinate | Sodium succinate hexahydrate | Organic acid | 270.14 | 4 |
| Citrate | Sodium citrate dihydrate | Organic acid | 294.1 | 6 |
| Acetate | Spdium acetate trihydrate | Organic acid | 136.08 | 2 |

**Table S2.** Table with respirofermenter taxa (Fermentative) or not capable of fermentation in aerobic conditions (Respirator). Members of the family Rhizobiaceae have been identified in one or the other category. To avoid overestimating one category or another, and because members of the Rhizobiaceae family were not abundant in our sugar samples, we excluded them from the analysis.

| Family | Fermentative | Reference |
|---|---|---|
| Enterobacteriaceae | Fermentative | Estrela et al. 2020 |
| Enterococcaceae | Fermentative | Estrela et al. 2020 |
| Pseudomonadaceae | Respirator | Estrela et al. 2020 |
| Burkholderiaceae | Respirator | Coyne, 2014 |
| Rhizobiaceae | Both | Encarnación et al., 1995; Fuhrer et al., 2005: Marcondes de Souza et al., 2014 |
| Moraxellaceae | Respirator | Estrela et al. 2020 |
| Xanthomonadaceae | Respirator | Estrela et al. 2020 |
| Lachnospiraceae | Fermentative | Estrela et al. 2020 |

**Table S3.** ANOVA results for the roles of source temperature (temp), carbon source of isolation (carbon), and functional group (fermenter) on the estimated maximum growth rate from thermal performance curves.

| | D.f. | Sum of Sq. | Mean Sq. | F-value | P-value |
|---|---|---|---|---|---|
| Temp | 4 | 0.057 | 0.014 | 3.07 | 0.017* |
| Carbon | 3 | 0.17 | 0.057 | 12.16 | < 0.0001** |
| Fermenter | 1 | 0.595 | 0.595 | 127.41 | < 0.0001** |
| Temp:Carbon | 12 | 0.065 | 0.005 | 1.15 | 0.319 |
| Temp:Fermenter | 4 | 0.185 | 0.046 | 9.91 | < 0.0001** |
| Carbon:Fermenter | 3 | 0.024 | 0.008 | 1.68 | 0.171 |
| Temp:Carbon:Fermenter | 11 | 0.087 | 0.008 | 1.7 | 0.073 |
| Residuals | 235 | 1.097 | 0.005 | | |

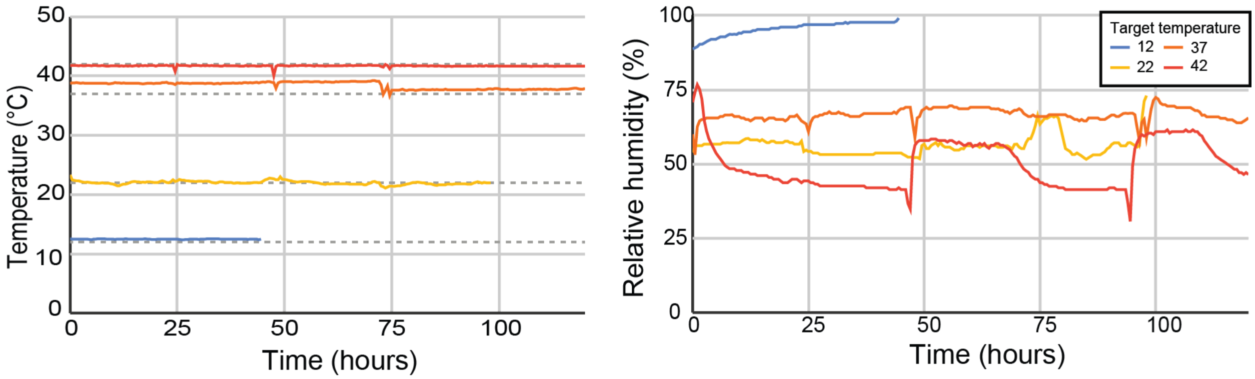


**Figure S1.** Treatment humidity and temperature conditions, over the first transfers. After the temperature and conditions settled we still periodically checked that conditions were maintained.

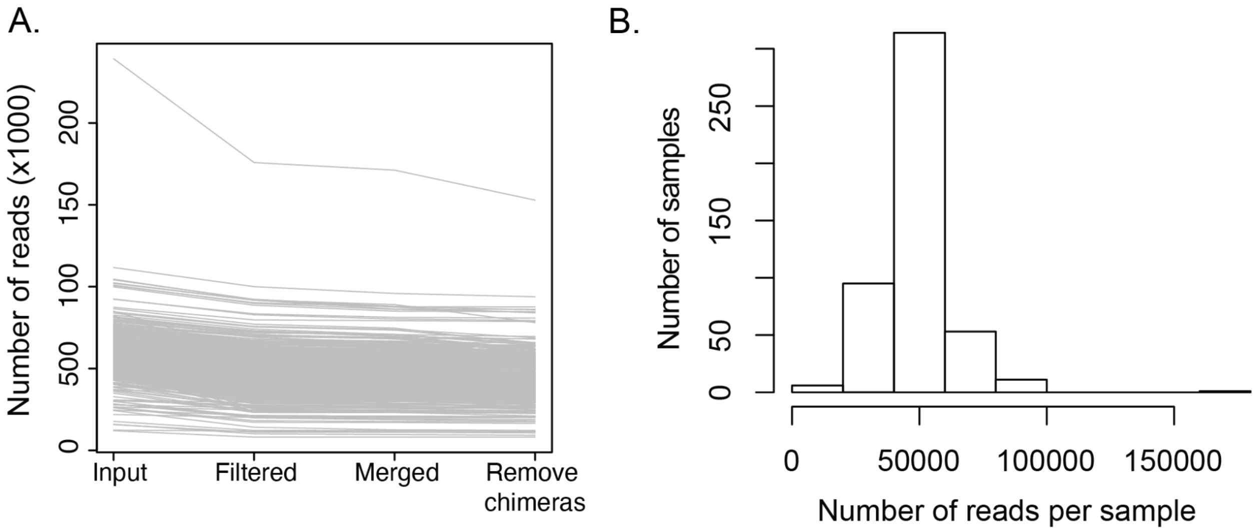


**Figure S2.** Most samples maintained a large proportion of reads throughout read processing (A). After cleaning, we still got a good coverage per sample with a median of 50,000 reads and a minimum of 8064.

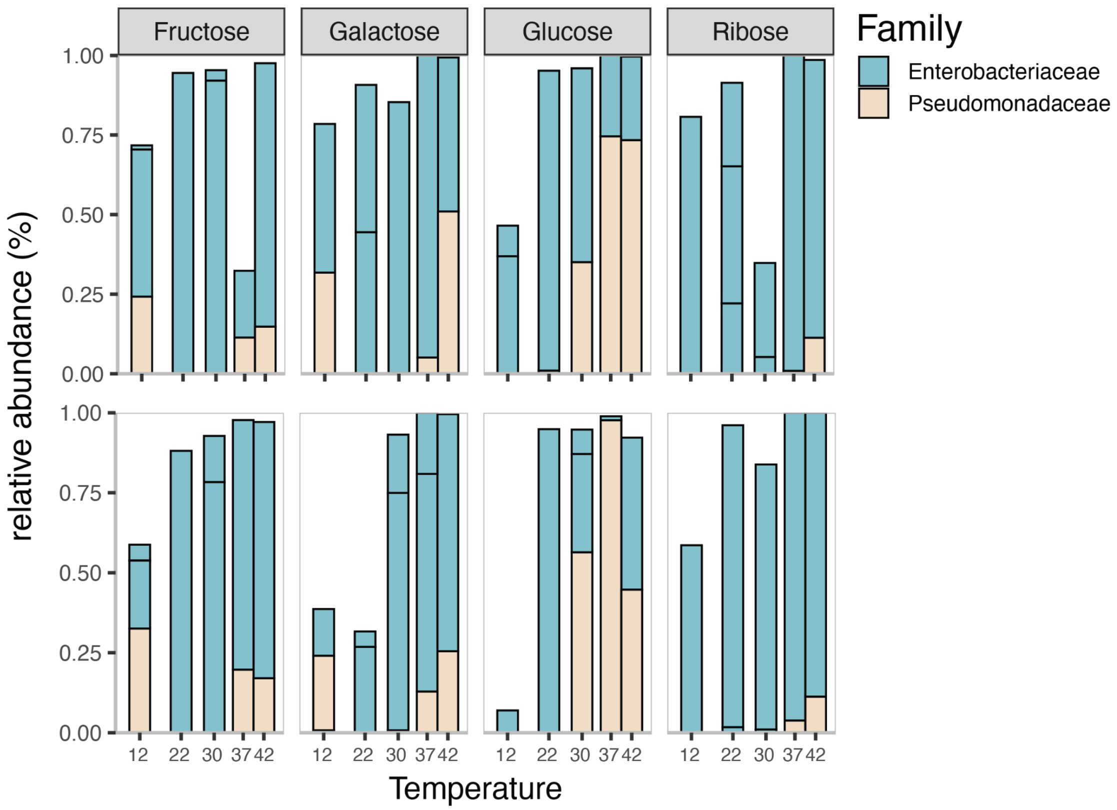


**Figure S3.** Mapping of isolates in 16S data (from Sanger sequences). Isolates are a good representation of the taxonomic composition of their source communities (the top panel is for the replicate community 2, and the bottom panel is for the replicate community 4).

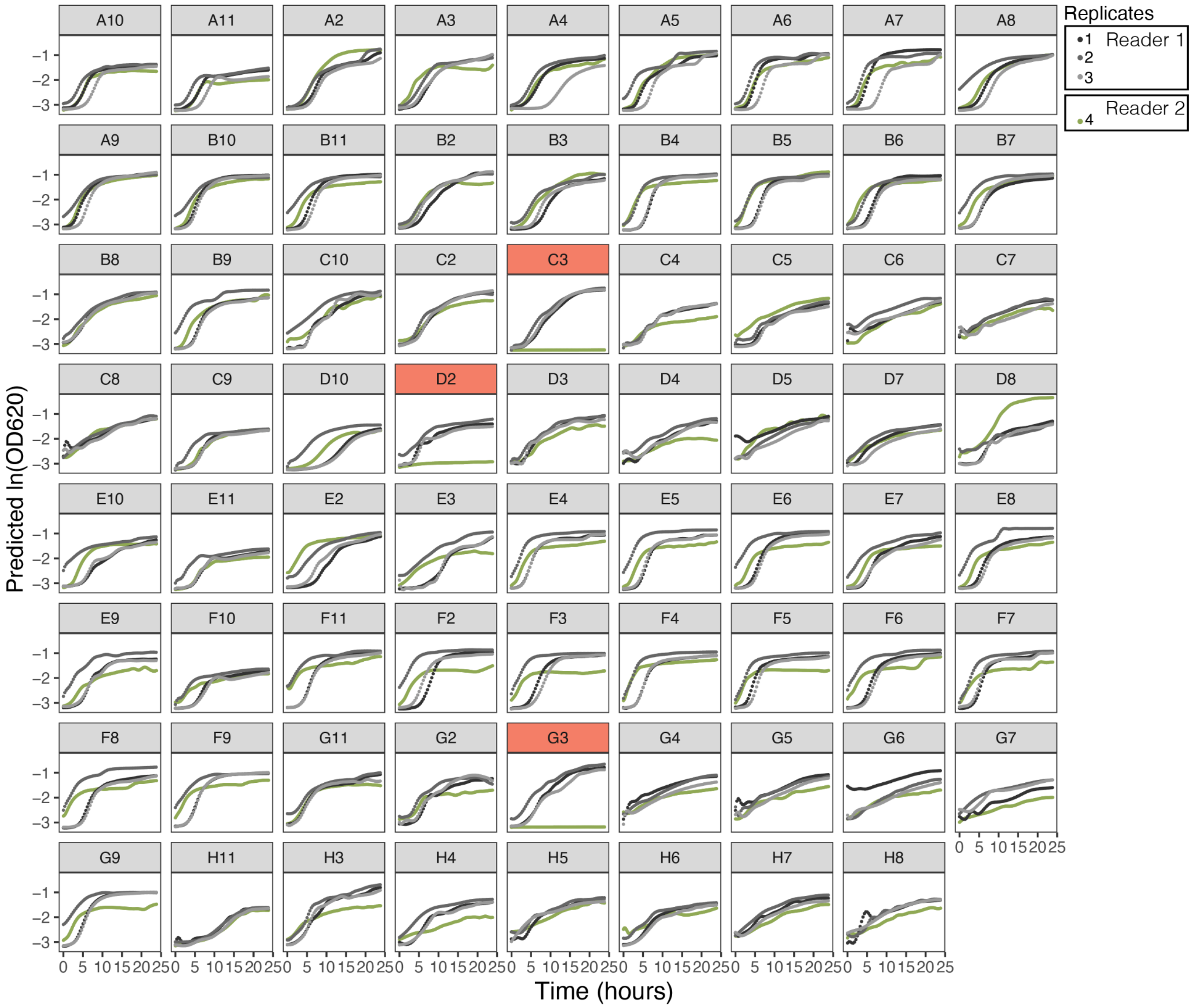


**Figure S4.** Growth curve fit of isolates in glucose at 30°C. Except for three isolates that did not grow (all originally isolatesd at 12°C) in the second batch, repliactes are consistent even across plate readers. The first three replicates were performed using a plate stacker together with a Biotek Epoch 2 in a room with controlled temperature set at 30°C, the fourth replicate was done much later after growing isolates again and using a Thermo accuSkan plate reader with internal control of temperature (set at 30°C).

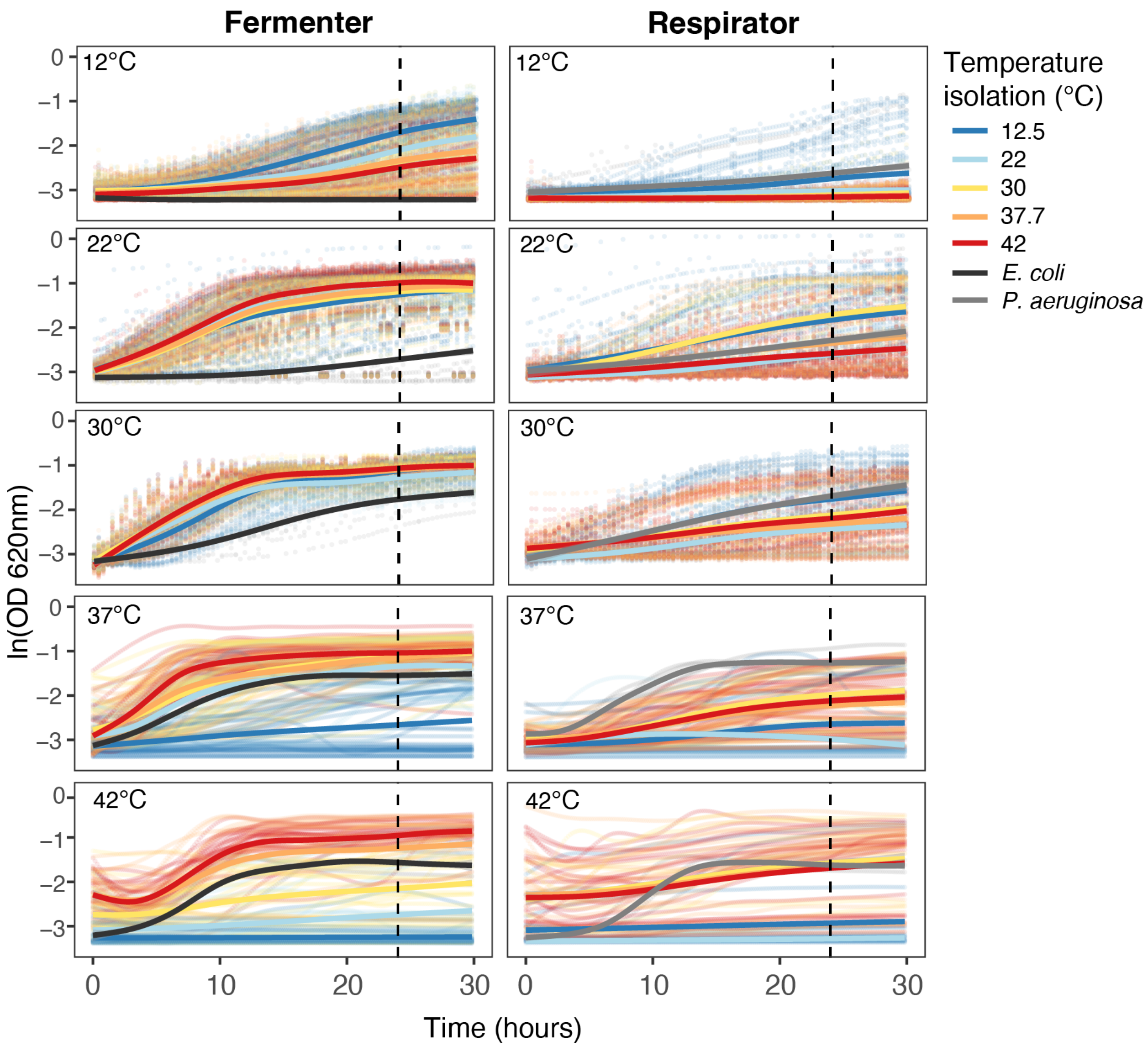


**Figure S5.** Growth curves of all isolates across different temperatures. Rows are the temperatures at which the growth curves were measured, columns separate isolates by functional group (fermenters, and respirators). *E. coli* and *P. putida* were used as reference strains for fermenters and respirators respectively. Except at 12°C, all isolates have reached stationary phase by 24hrs (vertical dashed line) when we sampled glucose consumption and organic acid production. Curves are coded according to the temperature of isolation of a particular strain. Points are curves fitted for each isolate and the lines are the means.

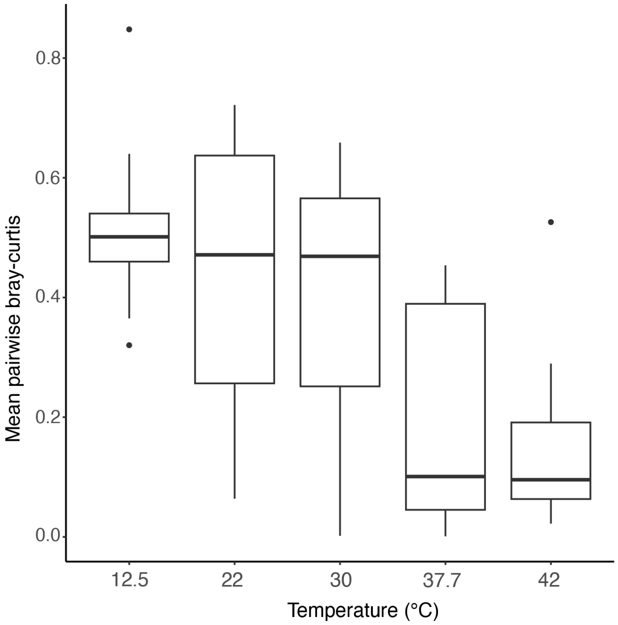


**Figure S6.** Mean pairwise Bray-Curtis between replicates within each carbon source, along different temperatures.

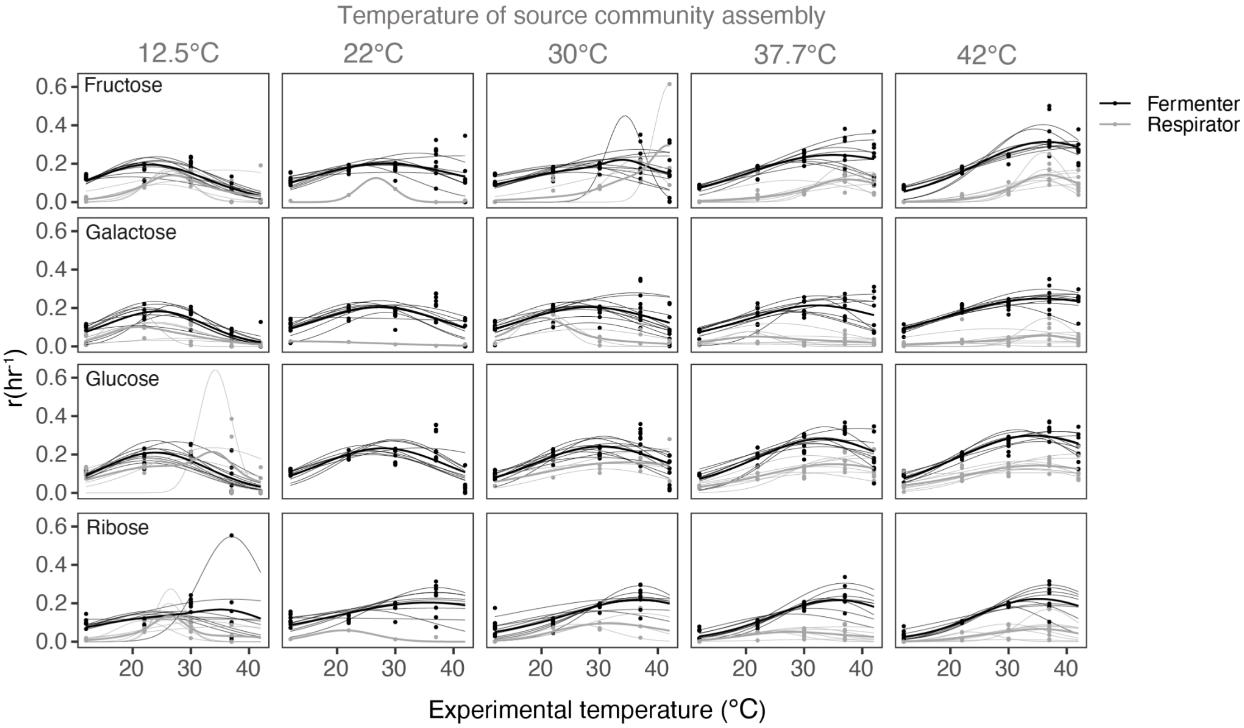


**Figure S7.** Response of growth rates of fermenters and respirators to increasing temperatures. A. Each column corresponds to a source temperature (the temperature at which the communities were assembled), and each row corresponds to the sugar used to measure growth.  Each thin line is the Gaussian model fit to one isolate, thick lines are the mean for fermenters (black) and respirators (gray).

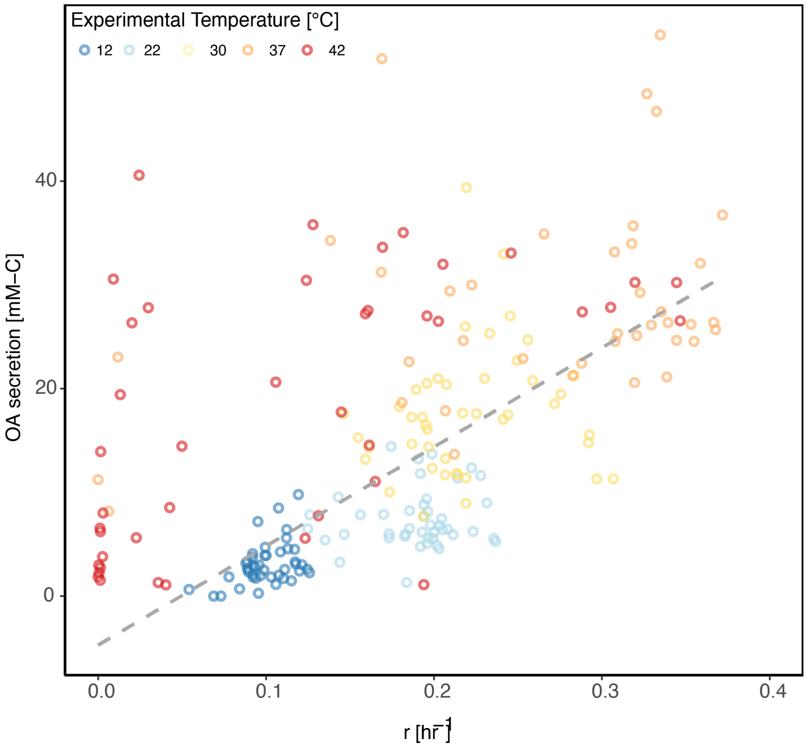


**Figure S8.** Total carbon that is secreted as organic acids (OA) at different growth rates, with points colored by their experimental temperature. Trendline shows linear fits to data excluding 42C.
